# Investigation of the pathogenic *RFC1* repeat expansion in a Canadian and a Brazilian ataxia cohort: identification of novel conformations

**DOI:** 10.1101/593871

**Authors:** Fulya Akçimen, Jay P. Ross, Cynthia V. Bourassa, Calwing Liao, Daniel Rochefort, Maria Thereza Drumond Gama, Marie-Josée Dicarie, Orlando G. Barsottini, Bernard Brais, José Luiz Pedroso, Patrick A. Dion, Guy A. Rouleau

## Abstract

A homozygous pentanucleotide expansion in the *RFC1* gene has been shown to be a common cause of late-onset ataxia. In the general population a total of four different repeat conformations have been observed: a wild type sequence AAAAG (11 repeats), and longer expansions of AAAAG, AAAGG and AAGGG sequences. However, in ataxia cases only the AAGGG expansion has been shown to be pathogenic. In this study, we assessed the prevalence and nature of *RFC1* repeat expansions in three adult-onset ataxia cohorts: Brazilian (n = 23) and Canadian (n = 26) cases that tested negative for other known ataxia mutations, as well as a cohort of randomly selected Canadian cases (n = 128) without regard to a genetic diagnosis. We identified the homozygous AAGGG pathogenic expansion in only one Brazilian family with two affected siblings, and in one Canadian case. The *RFC1* expansion may therefore not be a common cause of adult-onset ataxia in these populations. Interestingly we observed two new repeat motifs, AAGAG and AGAGG, which indicates the dynamic nature of the pentanucleotide expansion sequence. To assess the frequency of these two new repeat conformations in the general population we screened 163 healthy individuals. These novel motifs were more frequent in patients versus controls. While we cannot be certain that the homozygous genotypes of the novel expanded conformations are pathogenic, their occurrence should nonetheless be taken into consideration in future studies.

---

Autosomal recessive cerebellar ataxias are a heterogenous group of neurodegenerative diseases. Each type has distinct clinical characteristics, but the key symptom is progressive cerebellar dysfunction typically with gait and balance problems, dysarthria, dysmetria and oculomotor abnormalities. Other neurological dysfunction and/or non-neurologic phenotypes are also observed in some cases [1]. The most studied recessive ataxia is Friedreich’s ataxia (FRDA) which has the highest prevalence, followed by autosomal recessive spastic ataxia of Charlevoix-Saguenay (ARSACS), ataxia with vitamin E deficiency, autosomal recessive cerebellar ataxia type 1 (ARCA-1) and type 2 (ARCA-2), and ataxia with oculomotor apraxia type 1 (AOA-1) and type 2 (AOA-2) [2].

Homozygous expansions of an AAGGG pentanucleotide repeat in the second intron of the *RFC1* gene (hg19/GRCh37, chr4:39,350,045-39,350,103) were identified as a frequent cause of late-onset recessive ataxia, explaining the more than 20 percent of Caucasian sporadic ataxia patients [3]. A total of four distinct intronic repeat conformations were identified with different sequences: AAAAG_11_, as the wild-type sequence, and longer expansions of AAAAG_n_, AAAGG_n_ and AAGGG_n_. The configuration with the AAGGG pentanucleotide was shown to be the only disease-causing conformation of the expansion, ranging in size from 600 to 2,000 repeats.

*RFC1* represents a novel gene that could explain a significant number of adult-onset ataxia cases, the identification of cases in other populations may expand the clinical spectrum, provide example of variable regional prevalence and uncover repeat sequence differences. Therefore, we screened the *RFC1* expansion in Canadian and Brazilian ataxia patients.

Two cohorts consisting of adult-onset ataxia cases were used to estimate the prevalence of the *RFC1* expansions (Table 1). Cohort 1 and cohort 2 comprised Brazilian (n = 23) and Canadian (n = 26) adult-onset cases who did not carry variants in genes associated with common dominant and recessive ataxias (FRDA, DRPLA, SCA1, SCA2, SCA3, SCA6, SCA7, SCA10, SCA12, SCA17 and ARCA-1). Cohort 3 consisted of randomly selected adult-onset ataxia Canadian patients (n = 128). In addition, a cohort of 163 healthy Canadian control individuals was also examined to estimate the frequency of the novel sequence conformations that were observed for the *RFC1* repeat expansion. All subjects provided informed consent, and the study was approved by the appropriate institutional review boards.

**Table 1.** Allele frequency of *RFC1* repeat expansions in Brazilian and Canadian ataxia cohorts

|  | Cohort 1<br>n = 23 | Cohort 2<br>n = 128 | Cohort 3<br>n = 26 |
| --- | --- | --- | --- |
| Mean age at onset | 37 ± 7 | 50 ± 13 | 57 ± 10 |
| Prior genetic testing for<br>other common ataxias | + | - | + |
| AAAAG<sub>11</sub> | 29 (63%) | 193 (75.4%) | 36 (69.2%) |
| AAAAG<sub>n</sub> | 6 (13%) | 20 (7.8%) | 1 (1.9 %) |
| AAAGG<sub>n</sub> | 3 (6.5%) | 8 (3.1%) | 3 (5.8%) |
| AAGGG<sub>n</sub> | 5 (10.9%) | 16 (6.2%) | 11 (21.1%) |
| AAGAG<sub>n</sub> | 2 (4.3%) | 18 (7%) | 1 (1.9%) |
| AGGAG<sub>n</sub> | 1 (2.2%) | 1 (0.4%) |  |

Screening of the *RFC1* repeat expansion was performed on genomic DNA by RP-PCR as described in Cortese et al. [3]. RP-PCR products were separated on an ABI3730xl DNA Analyzer (Applied Biosystems®, McGill University and Genome Québec Innovation Centre) and results were visualized using GeneMapper® v.4.0 (Applied Biosystems®). The samples that were homozygous for the AAGGG repeat (according to the RP-PCR results) were subjected to long-range PCR (using the same primers as Cortese et al. [3]) and Sanger sequencing to examine the repeat sequence. Samples for which the allelic repeat combinations could not be determined by RP-PCR where subjected to a long-range PCR, the product of which was purified (QIAquick gel extraction kit, Qiagen). The Sanger sequencing results of these long-range PCR were analyzed using Unipro UGENE version 1.31 [4]. Finally, to test the association of the novel AAGAG variant with ataxia, we performed Fisher’s exact test using Canadian cases and controls.

To examine the prevalence of *RFC1*-based adult-onset ataxia, we screened the repeat expansions in a Brazilian and two Canadian cohorts. Based on the RP-PCR results of the Brazilian cohort, we had a total of four candidate patients that appeared to be homozygous for the pathogenic AAGGG expansion. However, Sanger sequencing revealed that two of these candidates were actually homozygous for the AAAGG expansion. Although different sets of primers were used in the RP-PCR, it seems that the AAAGG expansions can sometimes mimic AAGGG, misleading the results.

We identified three patients in the three cohorts with a homozygous pathogenic AAGGG repeat expansion. Two of the patients were Brazilian siblings and the other one was Canadian (Figure 1a and 1b respectively). Clinical features of the three patients with biallelic AAGGG expansions are summarized in Supplementary Table 1. The allele count and frequency of the different repeat expansions observed in all three cohorts are shown in Table 1.

**Figure 1.**
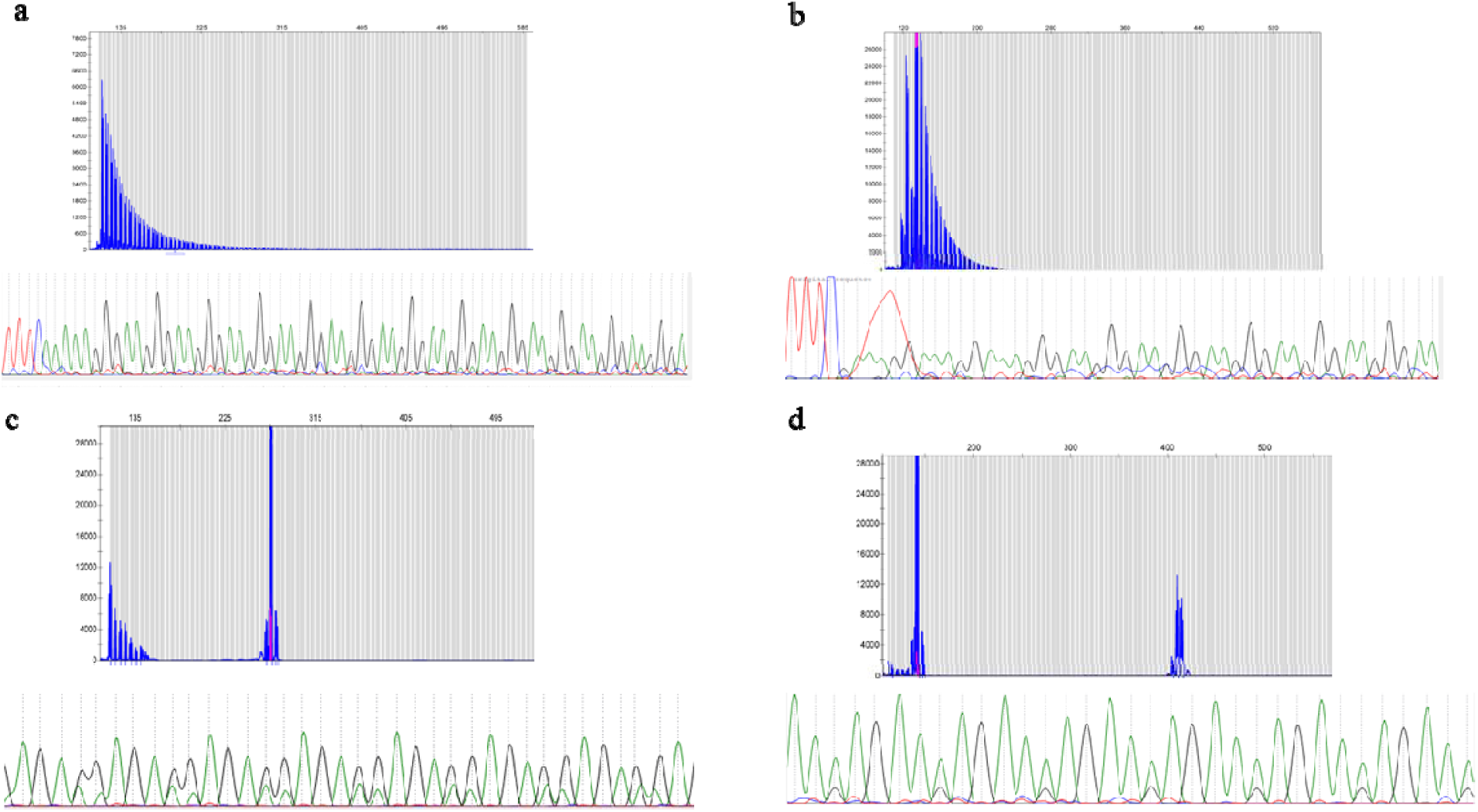
Repeat-primed PCR reactions targeting the AAGGG repeated motif. Fragment plots and Sanger chromatograms of long-range PCR results are shown for AAGGG-homozygous Canadian (1a) and Brazilian (1b) patients. Novel AGGAG (1c) and AAGAG (1d) expansion motifs (heterozygous) required both methods for identification.

Two new repeat expansion motifs were observed in our cohorts: AAGAG and AGAGG. The RP-PCR plots were characterized by a single peak, but the allele was longer than the wild type (Figure 1c and 1d). The novel motifs were in a heterozygous state in all 22 carrier individuals (Table 1). The average length of these novel expansions is 900 bp (180 repeats) (Supplementary Figure 1).

Additionally, we assessed the frequency of the AAGAG and AGAGG motifs in 163 healthy Canadian individuals using the same RP-PCR approach, followed again by long-range PCR and Sanger sequencing for samples where a novel expansion pattern was revealed by the RP-PCR. We detected a total of seven samples with a heterozygous AAGAG expansion confirmed by Sanger sequencing for an allele frequency of 2%. The frequency of the novel motif expansion was found to be approximately five times higher in cases than controls (Fisher’s exact test p = 2.95 × 10^−4^, OR = 5.10, 95% CI = 1.93 □ 15.16).

This study provides a detailed examination of the *RFC1* repeat region in Canadian and Brazilian cohorts with adult-onset ataxia. Two novel repeat motifs were identified. In addition to the cohorts in which known ataxia genes were ruled out, we used a Canadian cohort of adult-onset ataxia cases, without prior genetic testing, to estimate the frequency of *RFC1* pentanucleotide repeat motifs in this population. In contrast with the previous study where the *RFC1* AAGGG expansion explained a large proportion of familial and sporadic ataxia cases [3], the overall frequency of that *RFC1* repeat expansion mutation was very low in both our Canadian and Brazilian cohorts. The low prevalence of the *RFC1* pathogenic expansion, as well as identification of novel motifs might be due to the different genetic backgrounds of the Canadian and Brazilian populations [5]. Therefore, additional populations should be tested for the same pathogenic repeat, in order to draw a clear conclusion on its frequency in adult-onset ataxia.

The frequency of allelic configurations has been described only in healthy controls before [3]. This is the first study to assess the frequency of all possible configurations in a cohort of patients. In the previous study [3], a total of 3 % of the variants were not identified in the *RFC1* repeat loci in healthy cohort, and the existence of other possible allelic configurations was suggested. Our study identified novel repeat motifs in the *RFC1*, with a frequency of 0.07 and 0.02 in Canadian cases and controls respectively. The higher frequency of the novel AAGAG repeat unit in cases compared to controls suggests that it may also be associated with adult-onset ataxia. Further studies are warranted to confirm the structure and sequence of the repeated region, and to investigate potential biological impacts.

The presence of pathogenic or nonpathogenic repeat sequences in either disease-associated or wild type alleles has been observed across several expansion-associated diseases such as SCA37 [6, 7], SCA10 [8] and FRDA [9]. Variations interrupting the pure repeat sequences of disease-causing alleles can affect their penetrance, as well as the age at onset and severity of the conditions associated with specific repeats. Whereas interruptions in the normal alleles prevent the pathogenic expansions and provide the stability of repeats in disease-causing alleles. We did not observe the new sequence motifs along with a pathogenic AAGGG expansion in any of the patients, therefore further studies will be required to determine whether they affect the size of the pathogenic allele or the disease severity.

In conclusion, given the dynamic nature of the *RFC1* repeat, multiple validations of sequences and repeats length should be performed. To prevent false positive results, the RP-PCR plots should be interpreted with caution, and each AAGGG-positive sample should be validated by Sanger sequencing to confirm its true sequence. Additional work is needed to determine the frequency of other pentanucleotide repeat conformations and their association to adult-onset ataxia.

## Acknowledgements

We would like to thank the participants of the study. We thank Vessela Zaharieva and Hélène Catoire, for their assistance. Fulya Akçimen is supported by a Healthy Brains and Healthy Lives (HBHL) graduate student fellowship. GAR holds a Canada Research Chair and is funded by the CIHR.

