## Supplemental Data for "Investigation of the pathogenic *RFC1* repeat expansion in a Canadian and a Brazilian ataxia cohort: identification of novel conformations"

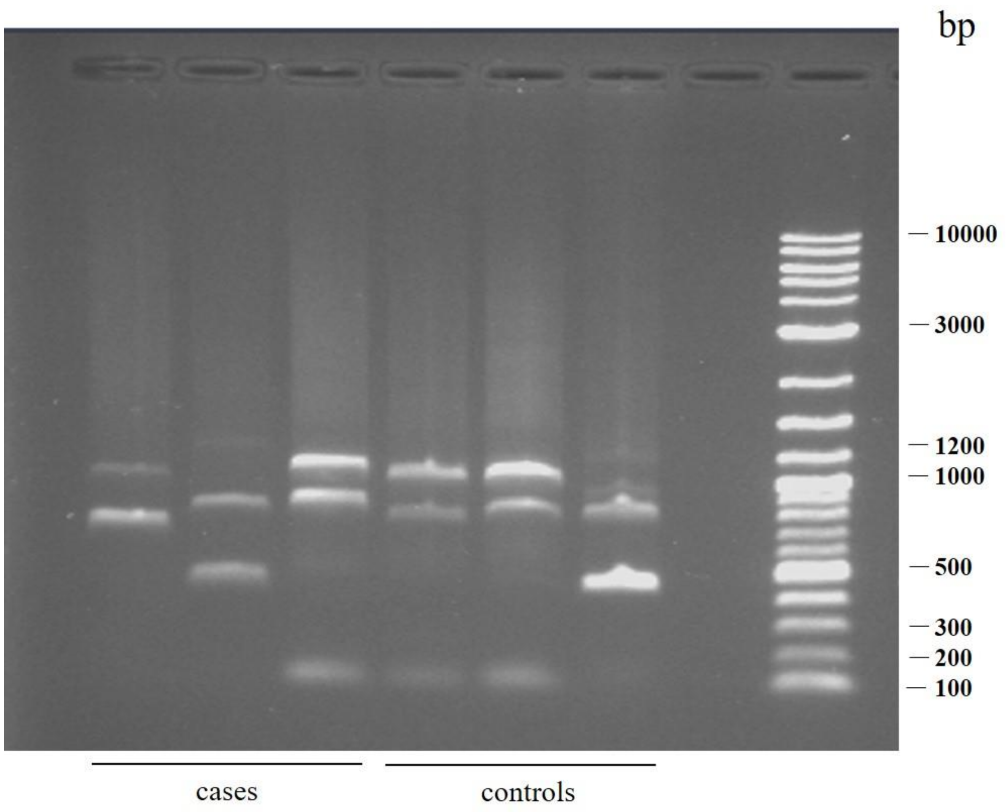


**Supplemental Figure 1**. Long-range PCR amplification shows the average length of the novel AGAAG repeats are 180 pentanucleotides.

**Supplemental Table 1.** Clinical features of patients carrying the recessive AAGGG repeat expansion in *RFC1*

| sample | origin | gender | family history | age at onset | age at examination | symptom at onset | neuropathy | cerebellar ataxia | nystagmus | cerebellar atrophy | SARA | other |
|---|---|---|---|---|---|---|---|---|---|---|---|---|
| Fam I-I | Brazilian | female | yes | 45 | 58 | Dizziness, gait and balance problems | sensorimotor axonal polyneuropathy | yes | yes | yes | 25 | Dysarthria, brisk tendon reflexes, vestibular areflexia |
| Fam I-II | Brazilian | female | yes | 45 | 56 | Dizziness, gait and balance problems | sensorimotor axonal polyneuropathy | yes | yes | yes | 27 | Dysarthria, brisk tendon reflexes, vestibular areflexia |
| Fam II-I | Italian | female | yes | 55 | 58 | Dizziness, gait and balance problems | None | yes | yes | yes | NA | Abnormal somatosensory evoked potentials, brisk tendon reflexes |

*SARA: Scale for the assessment and rating of ataxia.
